# Thalamic nuclei in patients with chronic facial pain: gray matter volume patterns before and after surgery

**DOI:** 10.1101/2023.12.20.572277

**Authors:** Pashkov Anton, Filimonova Elena, Zaitsev Boris, Martirosyan Azniv, Moysak Galina, Rzaev Jamil

## Abstract

Trigeminal neuralgia is a prevalent chronic pain disorder characterized by recurring episodes of intense facial pain, which significantly impairs patients’ quality of life. MRI-based biomarkers have consistently demonstrated their ability to predict pain intensity and treatment outcomes. However, most studies have primarily focused on the trigeminal system, paying less attention to the extensive neural reorganization that occurs throughout the brain in response to chronic pain. In this study, we aimed to examine the thalamus, a key brain structure involved in information processing, and provide a detailed perspective on thalamic remodeling in response to chronic pain at the level of individual thalamic nuclei. We analyzed a sample of 62 patients with primary trigeminal neuralgia undergoing surgical treatment, along with 28 healthy participants. Our results revealed significant gray matter volume changes in thalamic nuclei among patients with trigeminal neuralgia. Notably, the intralaminar nuclei (centromedian/parafascicular) and nuclei associated with visual and auditory signal processing (lateral and medial geniculate bodies) exhibited significant alterations, contrasting with the ventral group nuclei involved in nociceptive processing. Additionally, we found no substantial volume increase in any of the studied nuclei following successful surgical intervention 6 months later. The volumes of thalamic nuclei were negatively correlated with pain intensity and disease duration. The findings obtained in this study, albeit preliminary, have promising clinical implications as they unveil previously unknown facets of chronic pain development.

**Perspective:** The study examined alterations in gray matter volume within the thalamus of patients diagnosed with trigeminal neuralgia at the level of specific nuclei. The most significant changes were observed in the lateral and medial geniculate bodies, along with the pulvinar nuclei.

**Highlights:**

- A detailed investigation of the thalamic nuclei in patients suffering from primary trigeminal neuralgia (TN) has been carried out for the first time.
- Patients with TN showed a significant gray matter volume decrease in intralaminar, sensory and associative nuclei.
- Following successful surgery, there was no observed increase in the volume of the investigated nuclei.

## Introduction

Trigeminal neuralgia is a prevalent chronic pain disorder characterized by intermittent attacks of intense electric shock-like facial pain, ranging from a few seconds to several minutes in duration. The sharp and severe pain experienced by patients with classical trigeminal neuralgia is commonly attributed to vascular compression of the trigeminal nerve root, with the superior cerebellar artery being the most frequently implicated offending vessel [27]. Many patients with trigeminal neuralgia can effectively manage their pain by using tailored doses of antiepileptic drugs, tricyclic antidepressants, or combined norepinephrine-serotonin reuptake inhibitors, which have demonstrated clinical efficacy and relative safety in pain management [17]. However, some patients may experience a loss of effectiveness of the prescribed medications in controlling the frequency or intensity of pain attacks over time. Although there are non-surgical therapeutic alternatives available, such as lidocaine and botulinum toxin injections or the implementation of transcranial magnetic stimulation protocols targeting specific brain regions like somatosensory, motor, and dorsolateral prefrontal cortices [48], surgical treatment remains one of the most effective approaches for alleviating trigeminal pain [29].

In clinical practice, modern neuroimaging protocols are routinely used to assess the presence and severity of neurovascular conflict and evaluate microstructural changes in the trigeminal nerve [47]. MRI-derived biomarkers have repeatedly demonstrated their ability to predict pain intensity and treatment outcomes [23]. However, most of these studies have focused solely on the trigeminal system, overlooking the fact that chronic pain leads to extensive neural reorganization throughout the brain [18,30]. Among the brain regions affected by prolonged pain experiences, the thalamus is frequently highlighted for its prominent role in sensory computations, including the processing of nociceptive signals. Recent publications have provided evidence of significant gray matter volume (GMV) reduction in the thalami of patients with trigeminal neuralgia [11,33,38,46]. Furthermore, the thalamus is increasingly recognized as a crucial hub not only for relaying neural signals to upstream areas but also for flexibly integrating diverse channels of incoming information based on the current environmental context and internal states [24,37]. Considering this perspective, a more nuanced understanding of thalamic involvement is necessary to comprehend both acute and chronic pain-related behavior.

However, previous studies examining changes in thalamic GMV have often treated the thalamus as a single entity, without delving into specific structural alterations at the level of individual nuclei. In our study, we aimed to advance this understanding by taking a more detailed approach and dividing the thalamus into 25 distinct nuclei. Our objective was to provide a finer-grained perspective on the remodeling of the thalamus in response to chronic pain, additionally focusing on the associations between gray matter alterations and patients’ clinical data. Finally, we aimed to investigate whether these changes could be reversed following successful surgery.

## Materials and methods

### Patients and study design

This prospective study was conducted at our hospital from January 2022 to September 2023, and constituted a part of an ongoing scientific project dedicated to the comprehensive investigation of neuroimaging, neuropsychological, and blood biomarkers of chronic facial pain, as well as the search for predictors of surgical outcomes. The study included a total of 111 participants. Among them, there were 75 patients with a confirmed diagnosis of trigeminal neuralgia who were referred for surgical treatment due to the ineffectiveness or severe side effects of their pharmacological therapy (aged 27-83, 61% female). Additionally, a small number of patients (n=6) who were being managed conservatively on an outpatient basis at our Center were also included (aged 43-72, 67% female). Gender and age-matched healthy controls (n=30, aged 41-74, 57% female; HC) were recruited through local advertisements targeting hospital personnel.

The inclusion criteria for patients in this study were as follows: experiencing constant, episodic unilateral or bilateral pain in the innervation zone of the first and/or second and/or third branches of the trigeminal nerve; no contraindications for MRI; absence of intracranial structural anomalies and facial area anomalies (such as tumors, vascular malformations, and aneurysms) observed in neuroimaging data; absence of other neurological disorders (particularly migraines) and psychiatric disorders; MMSE score > 24 and right-handedness based on manual asymmetry profile. All patients with classical and idiopathic trigeminal neuralgia included in the study met the diagnostic criteria outlined in points 13.1.1.1 (Classical trigeminal neuralgia) and 13.1.1.3 (Idiopathic trigeminal neuralgia) of The International Classification of Headache Disorders, 3rd edition, 2018 (collectively referred to as primary trigeminal neuralgia, PTN). To minimize confounding factors, we specifically enrolled patients who met the criteria for these two nosological groups, excluding cases of trigeminal neuralgia secondary to multiple sclerosis, brain tumors, and other intracranial anomalies from consideration.

All selected patients and healthy controls underwent a series of diagnostic procedures, including brain MRI, as well as comprehensive neuropsychological and psychological assessment (not reported in this manuscript). Patients were additionally administered a number of clinical scales to estimate the levels of pain severity and potential postsurgical complications. The flowchart illustrating the patient selection process is depicted in Figure 1. Follow-up examination using the same protocol was conducted 6 months post-surgery.

**Figure 1.**
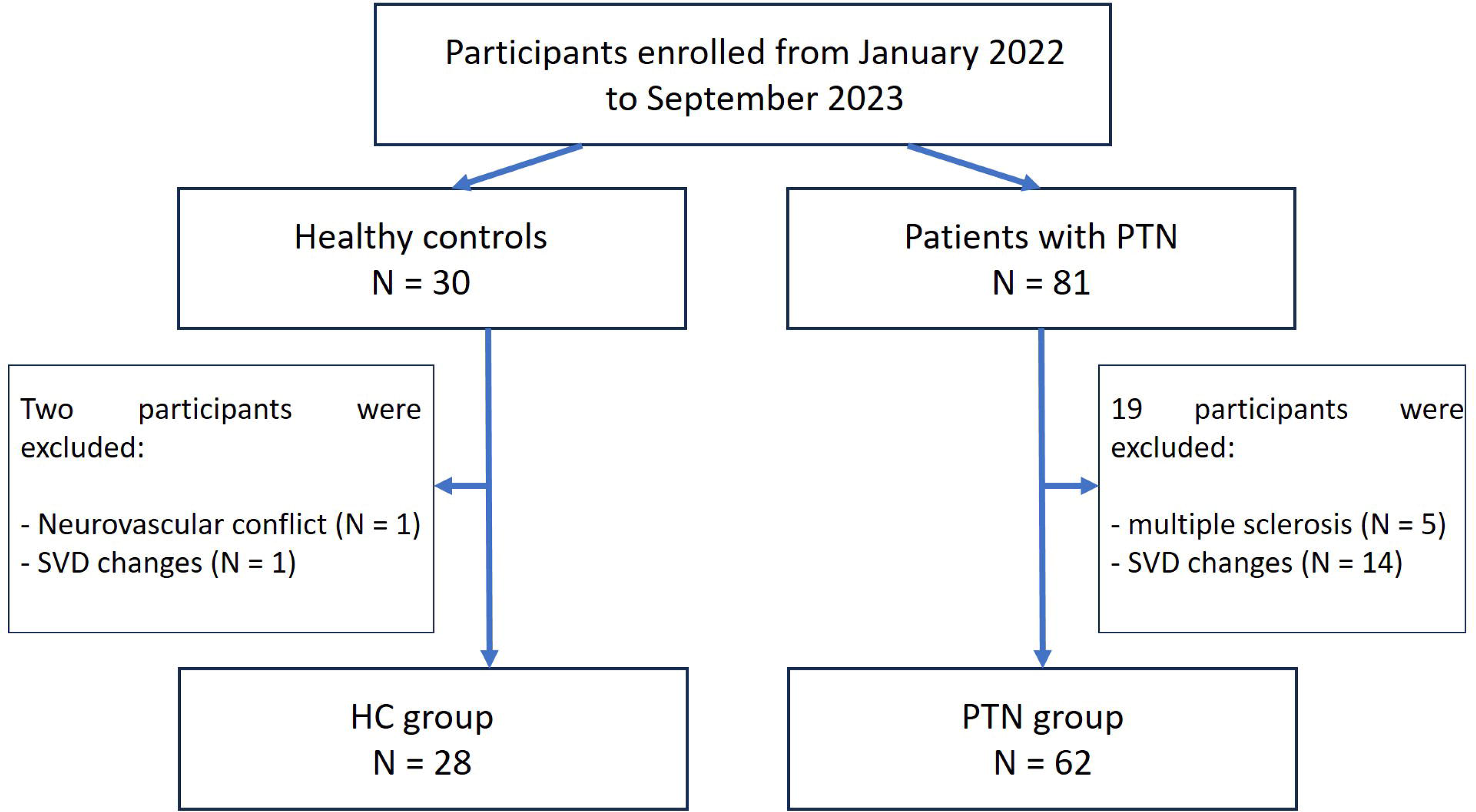
The flowchart of the current study.

All patients and healthy controls provided written informed consent to participate in the study, in accordance with the Declaration of Helsinki. Ethics approval for the research protocol was given by local ethic committee of Federal Center of Neurosurgery dated 25.05.21 (protocol # 7)

### MRI acquisition

MR imaging data were acquired using a 3T system (Ingenia, Philips Healthcare, The Netherlands) equipped with a 16-channel receiver head coil. Brain magnetic resonance imaging included conventional sequences, as well as high-resolution 3D T1-WI (for morphometry) and high-resolution 3D T2-WI FFE with axial orientation (for detection and classification of neurovascular conflict). The total acquisition time was approximately 30 minutes. The T1-WI high-resolution sequence (3D TFE in the sagittal plane) had the following parameters: TR -– 6.56 ms, TE – 2.95 ms, FOV – 256*256 mm, flip angle - 8, matrix – 256*256, and slice thickness – 1 mm.

### MRI data processing

Initially, conventional MRI data were analyzed by a neuroradiologist to find additional pathology not related to disease (such as prominent signs of small vessel disease, for example). Furthermore, cerebellopontine angle structures were visually evaluated with the 3D T2 FFE sequence in each case for the assessment of the presence and severity of neurovascular conflict, according to Sindou classification, as previously described [28].

Fully automatic surface-based MR morphometry of high-resolution T1-WI was performed with FreeSurfer v7.2.0 software (https://surfer.nmr.mgh.harvard.edu). Segmentation of the thalamic nuclei was performed as part of the FreeSurfer 7.2.0 functional. The results were visually checked by the neuroradiologist for each subject. The Figure 2 depicts thalamic nuclei segmented according to the protocol described in [25].

**Figure 2.**
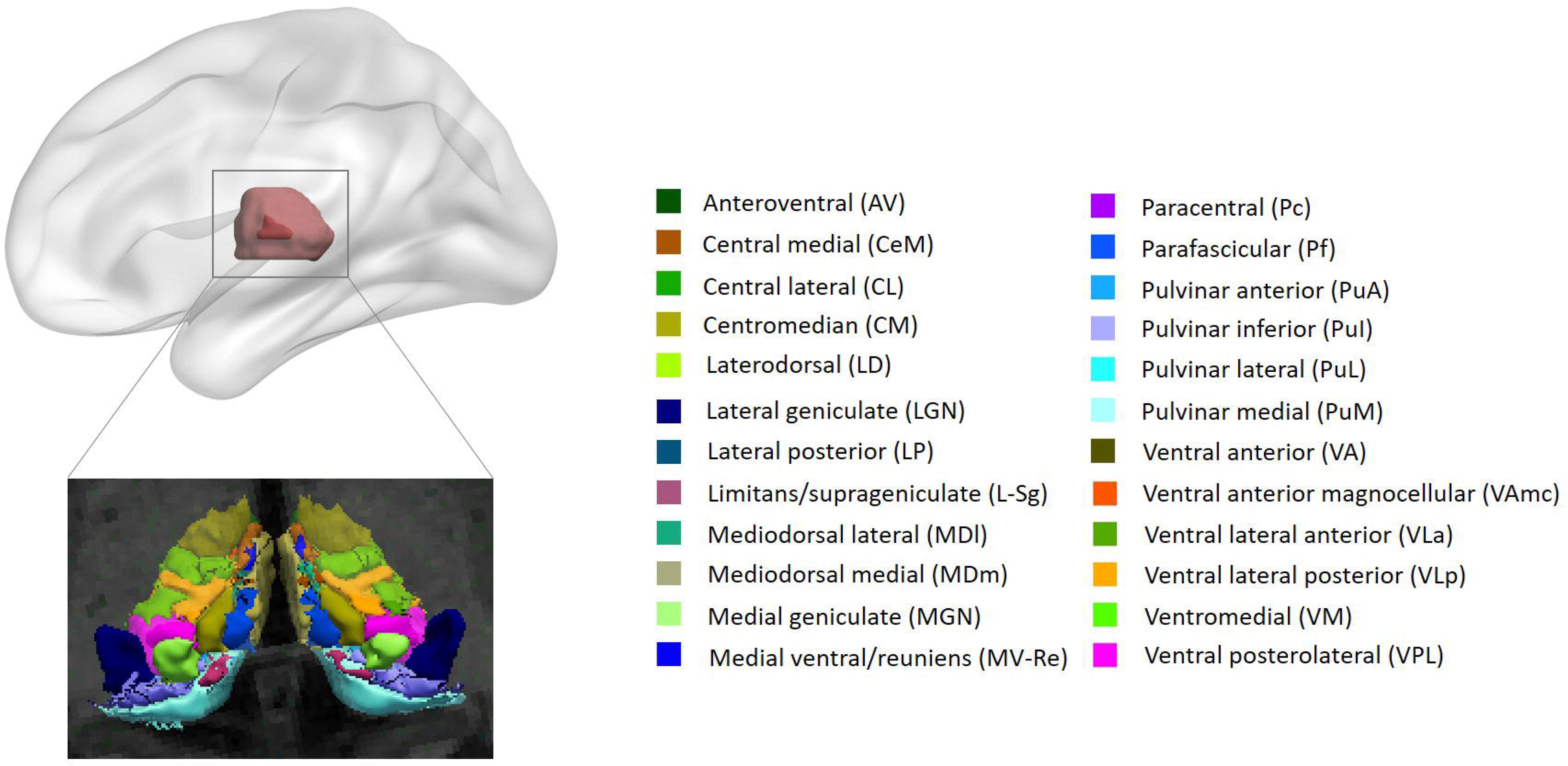
The example of thalamus segmentation into 25 separate nuclei on each side. The paratenial nucleus (Pt) is not shown due to its small size.

### Pain assessment

To assess the severity of a patient’s pain upon admission to the hospital and at the 6-month follow-up, two primary assessment tools were employed. The Brief Pain Inventory (BPI) utilizes a 10-point numerical rating scale and prompts patients to rate their minimal, maximal, and average pain intensity over the past week. Furthermore, the BPI evaluates the impact of pain on various aspects of the patient’s daily activities and aids in distinguishing clinically significant pain features, such as tingling, burning, or numbness sensations.

In addition, the Barrow Neurological Institute Pain Intensity Score (BNIPS) assigns grades ranging from I to V, providing valuable insights into the medications used by patients and their effectiveness in pain management. Grade I is assigned to patients who report no pain. Grade II is designated for patients experiencing occasional pain that does not require medication. Patients requiring prescribed medication for pain are classified as Grade III. Grade IV indicates inadequate pain control despite medication use. Finally, Grade V encompasses severe pain or lack of pain relief. For further analysis in this study, we exclusively utilized the mean BPI score.

### Statistical analysis

Continuous variables were reported as mean ± standard deviation if normally distributed and median (interquartile range, IQR) if distribution was different from normal. Normality of the distribution was checked with Shapiro-Wilk test. Categorical variables were presented with frequency and percentage. Chi-squared test was used for categorical data analysis. t-test with Welch correction was utilized for normally distributed data. Alternatively, we employed Mann-Whitney test when the normality assumption was violated. Comparison of measured variables in pre- vs post-surgery conditions in the same patients (paired sample) were done with Wilcoxon signed-rank test. In addition, homogeneity of variances in experimental groups were checked using a Levene test. We divided the group of patients with trigeminal neuralgia into two subgroups: those with right-sided pain and those with left-sided pain. Each subgroup was compared to a control group. One-way ANCOVA with post hoc Tukey tests was used to compare mean values of gray matter volumes between groups while controlling for age, gender and intracranial volume variables. Generalized eta squared 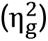 was utilized as a measure of effect size. To assess a relationship between brain and clinical variables Spearman correlations were employed. p-value of 0.05 was considered a threshold for evaluating statistically significant results. False discovery rate (FDR) correction was applied to adjust the p-values under multiple comparison conditions. All p-values reported in the main text were FDR-adjusted. All statistical analyses were run in R (v. 4.3.1, 2023).

## Results

### Demographic and clinical data

Total of 62 patients (33 female) and 28 healthy controls (16 female) were considered for the final analysis. Out of the initial sample of 81 patients, five participants were excluded due to multiple sclerosis and 14 patients had significant white matter lesions (Fazekas grades 2 or 3). Two patients were excluded from the control group: one patient was found to have a neurovascular conflict on MRI not accompanied by facial pain, while the other exhibited white matter changes consistent with Fazekas grade 2.

The mean age of the recruited patients was 59.7±10.6 years. Patients and healthy controls did not differ in age (p > 0.1). Similarly, PTN patients did not differ from the control group in gender (p > 0.1) and body mass index (p > 0.05). However, the statistically significant difference between groups in education years was identified (U = 519.5, p = 0.008). Thirty seven patients (59.7%) had right-sided pain, twenty four patients (38.7%) suffered from left-sided pain and in one patient (1.6%) the pain was bilateral. Median disease duration for the selected patients was 8 years (IQR = 6.5). This data is summarized in Table 1. Overwhelming majority of patients had BNIPS grades IV or V (89%). Median value of BPI average scores were 5 (IQR = 3) and this value increased up to 8 (IQR = 4) for the BPI max scores. Before surgery, the most commonly taken drug was carbamazepine (95%), followed by gabapentin (16%), amitriptyline (8%) and pregabalin (5%). Neurovascular conflict of different grades was identified in 50 patients (81%). Fourteen patients (23%) had previously undergone surgery for trigeminal neuralgia and were admitted to our neurosurgical hospital due to pain recurrence. This group of patients did not differ from those undergoing surgery for the first time in terms of pain intensity (p > 0.05) and disease duration (p > 0.1). Similarly, patients undergoing surgical treatment in the hospital and patients receiving outpatient treatment did not differ in terms of pain intensity (p > 0.1) and disease duration (p > 0.1).

**Table 1.**
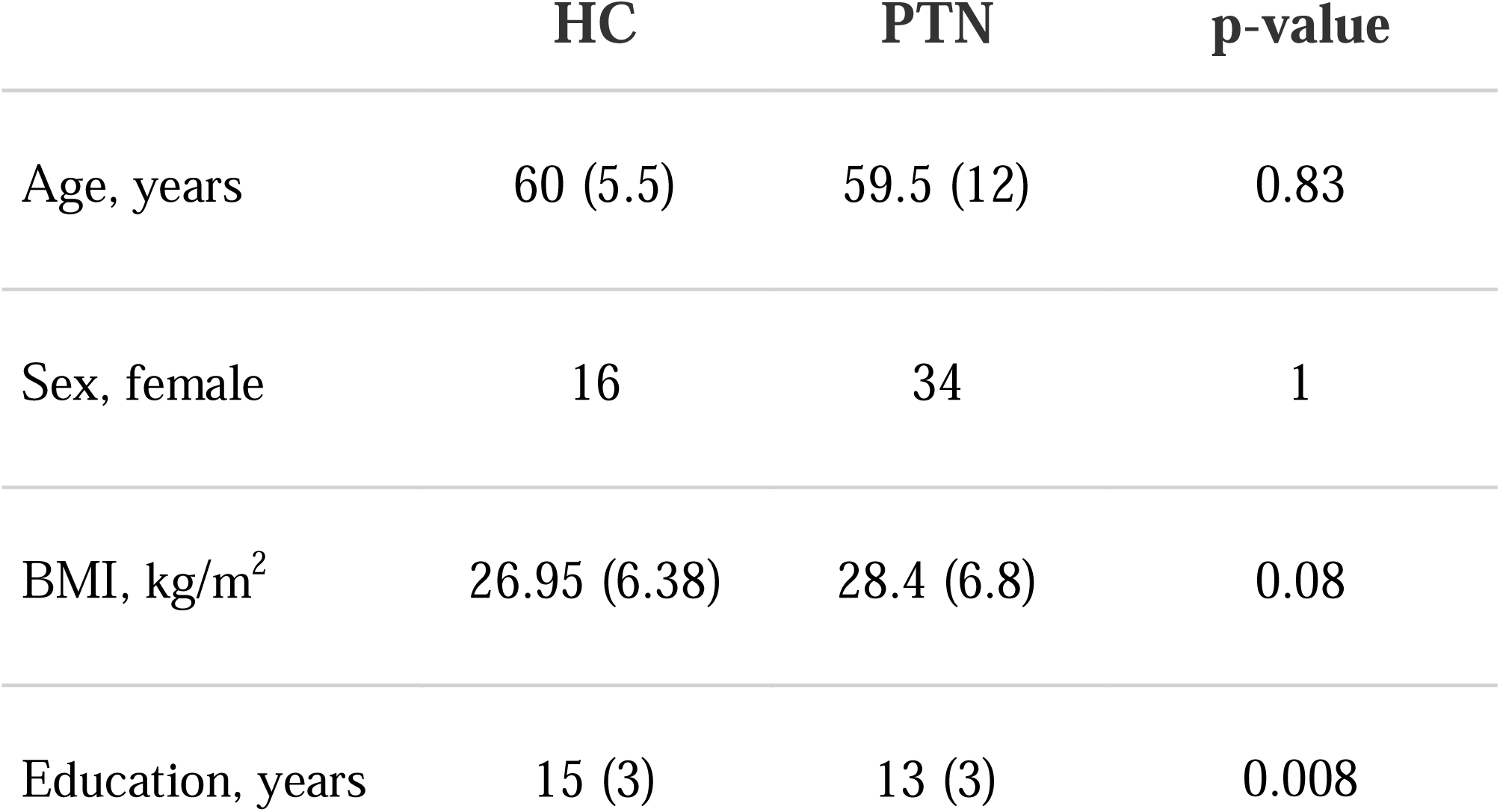
Comparison of basic demographic and health-related variables between experimental and control groups.

|  | HC | PTN | p-value |
| --- | --- | --- | --- |
| Age, years | 60 (5.5) | 59.5 (12) | 0.83 |
| Sex, female | 16 | 34 | 1 |
| BMI, kg/m<sup>2</sup> | 26.95 (6.38) | 28.4 (6.8) | 0.08 |
| Education, years | 15 (3) | 13 (3) | 0.008 |

**Table 2.**
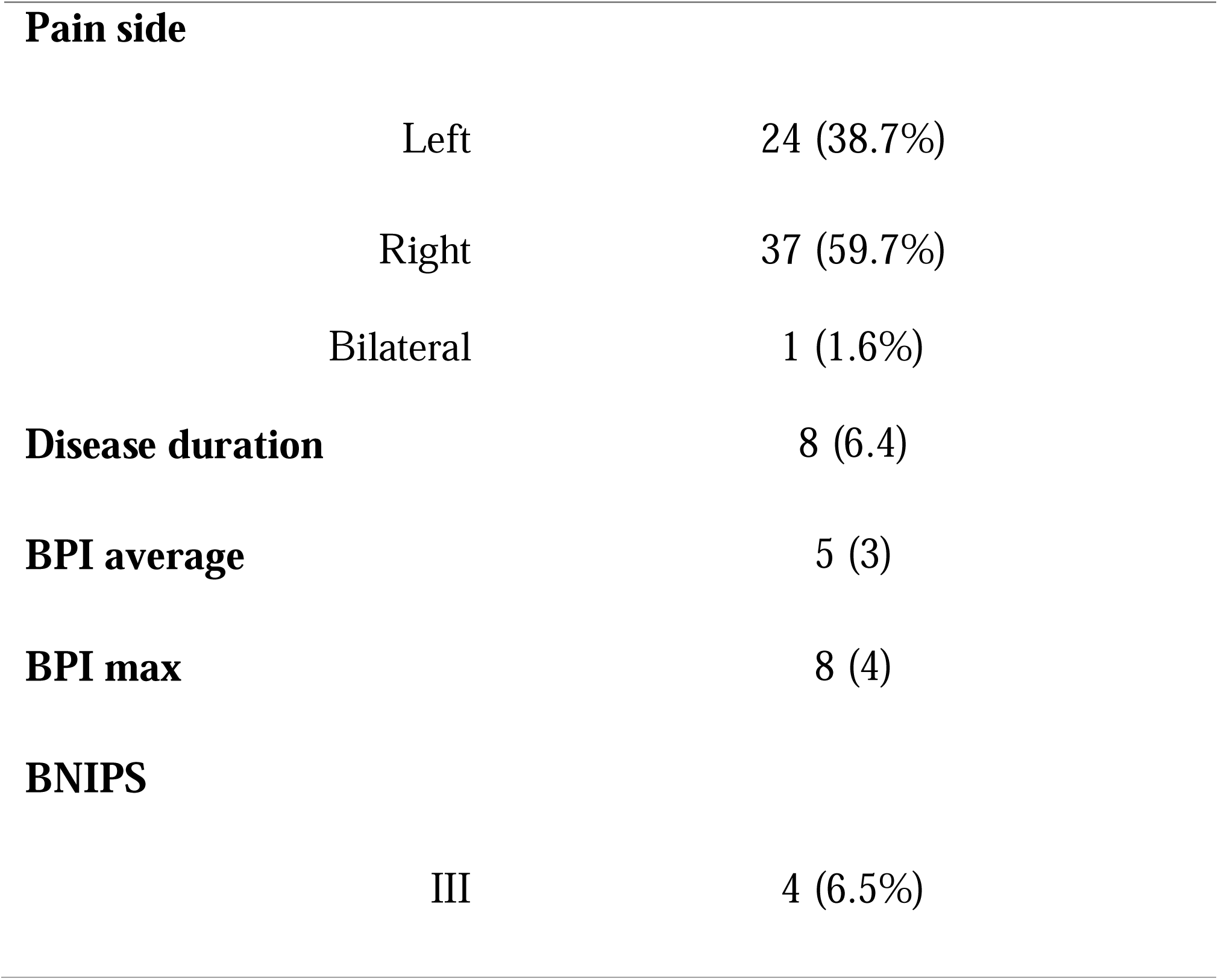
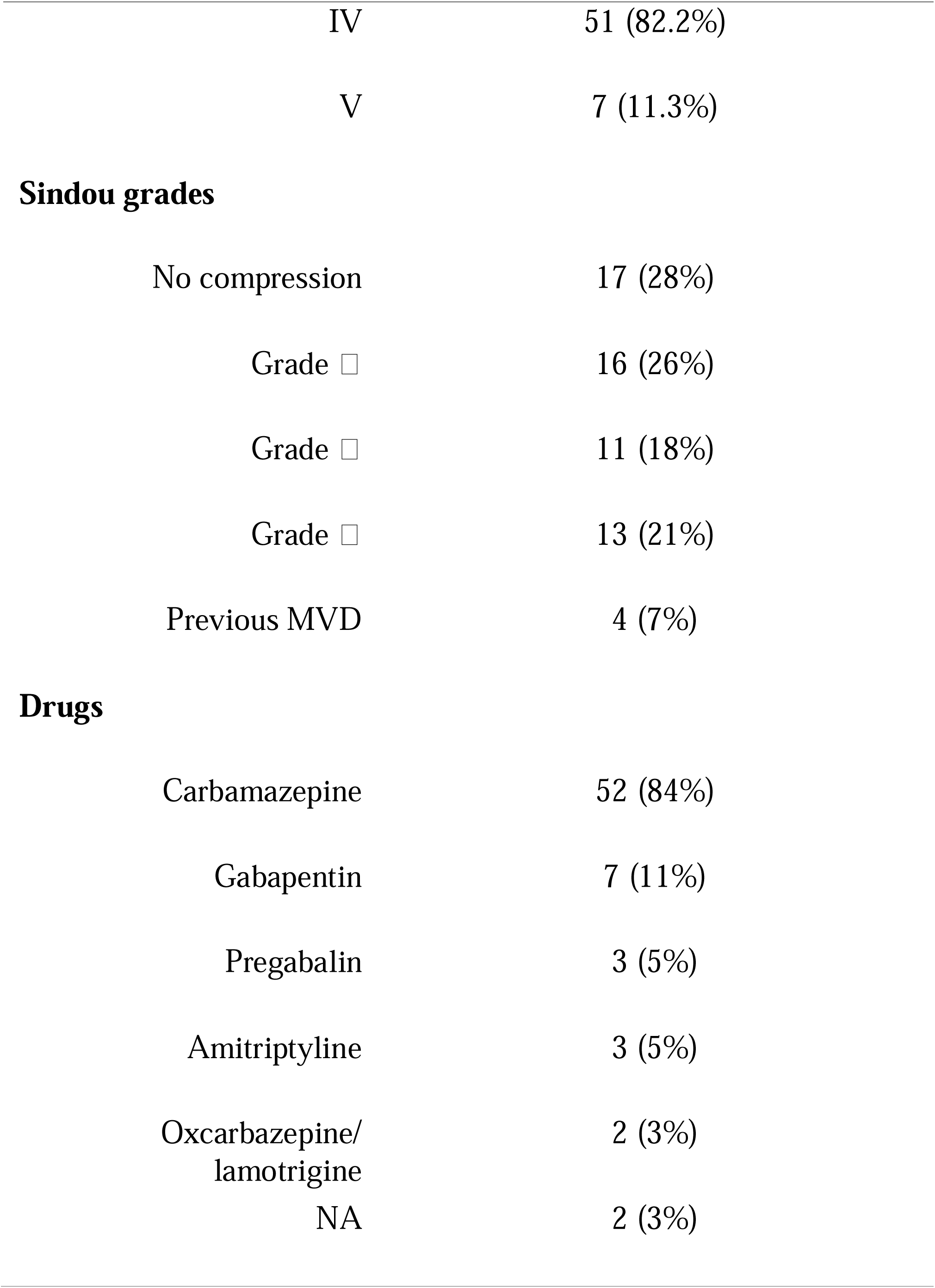
Major clinical variables in patients with PTN.

### Grey matter volume alterations in thalamic nuclei

We used one-way ANCOVA with post hoc Tukey tests and FDR-correction for multiple comparisons to estimate GMV differences in 25 thalamic nuclei on each side between TN patients and healthy controls (Table 3). Age, gender and intracranial volume variables were taken as covariates in the analysis. We revealed three clusters of nuclei showing significant differences between groups. First cluster consisted of intralaminar nuclei such as centromedian (CM), parafascicular (Pf) and paratenial (Pt) nuclei. Second cluster included lateral (LGN) and medial geniculate bodies (MGN) as well as thalamic nuclei belonging to the ventral group, mainly related to sensory processing. Third cluster was composed of so-called associative thalamic nuclei, namely, mediodorsal lateral (MDl), laterodorsal (LD) and pulvinar nuclei.

**Table 3.**
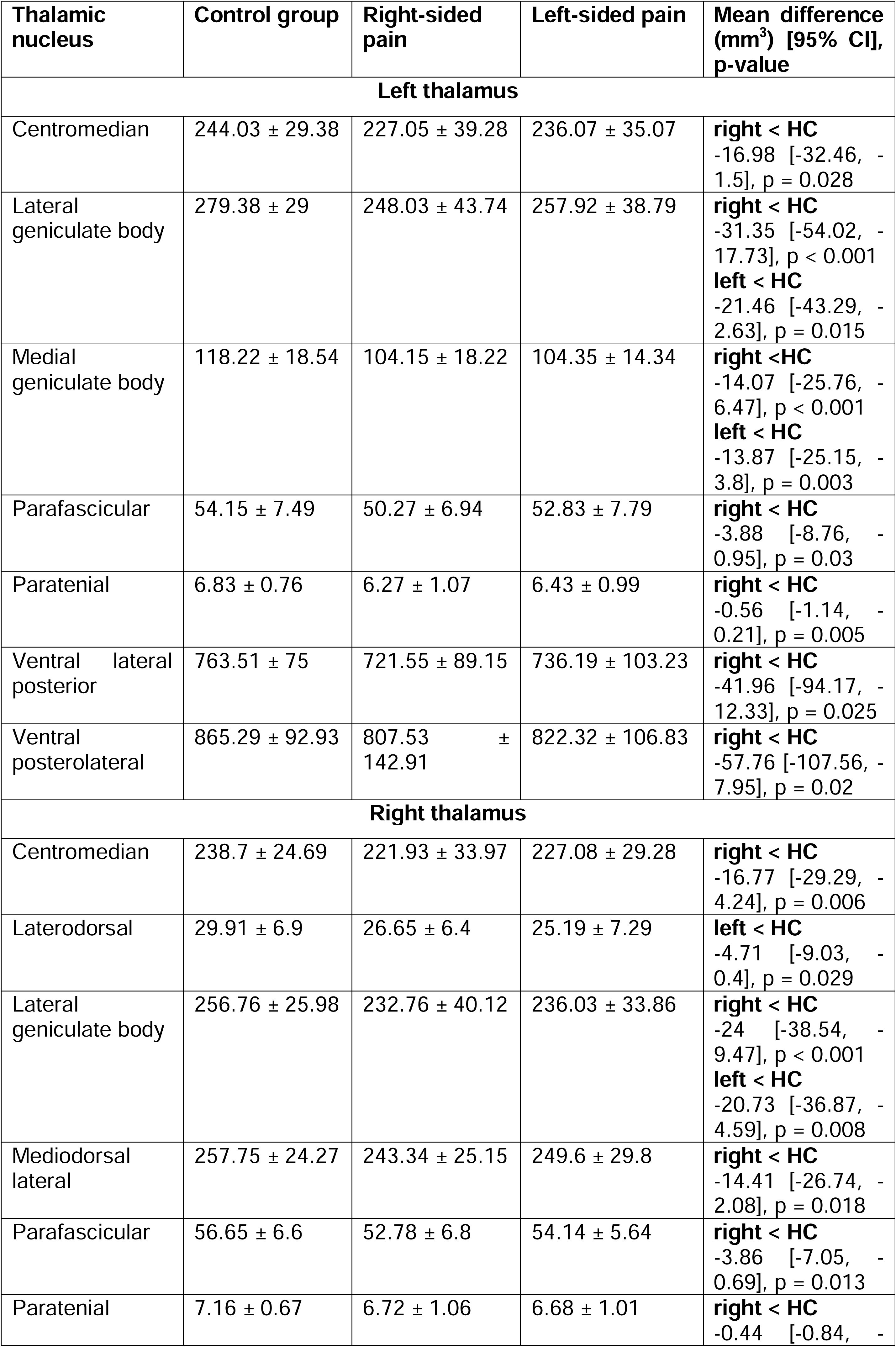
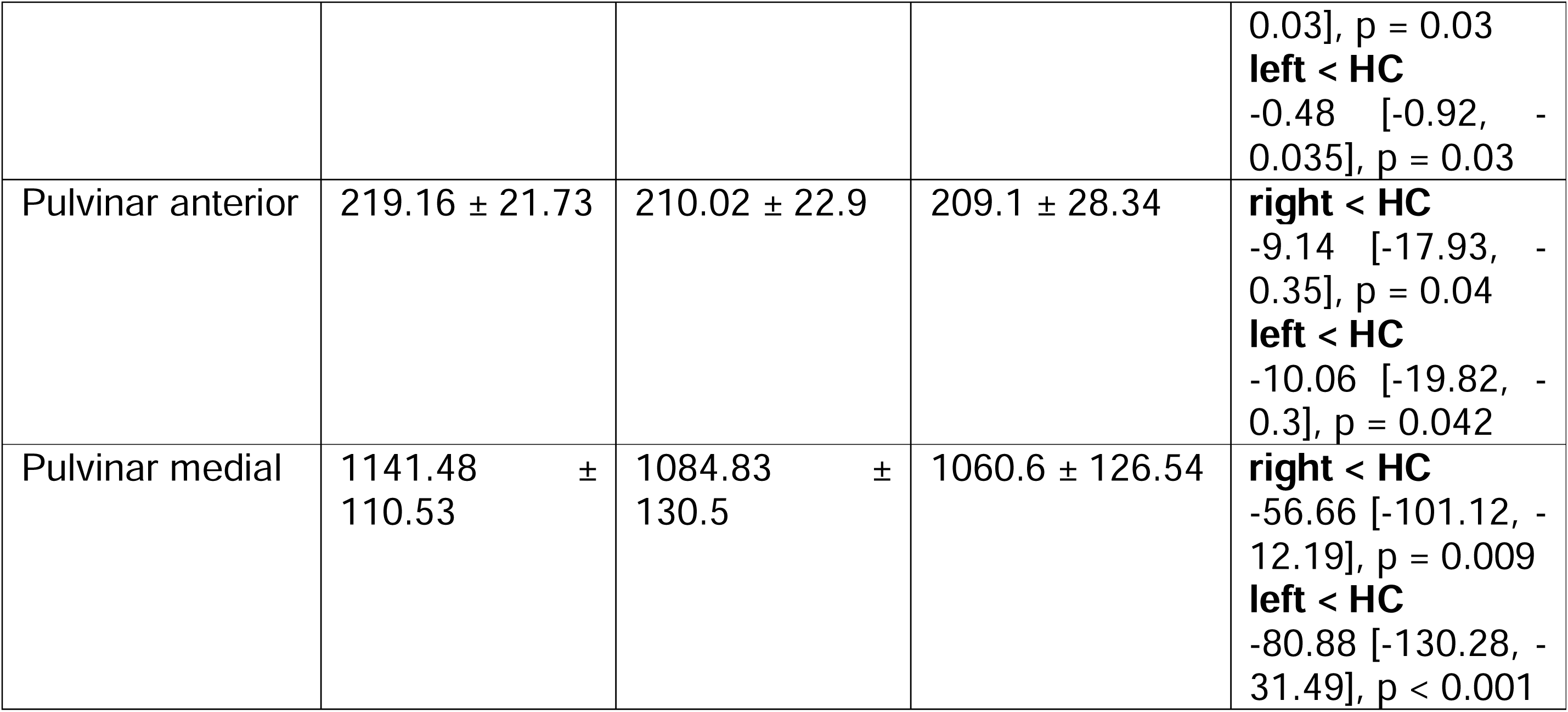
The gray matter volume of thalamic nuclei, demonstrating statistically significant differences between groups of healthy participants and patients with left- and right-sided pain.

Patients and healthy volunteers differed significantly in gray matter volumes in the left centromedian (F(2,83) = 3.45, p = 0.036, 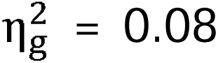) and parafascicular nuclei (F(2,82) = 3.49, p = 0.035, 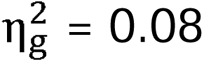). Patients with right-sided pain had smaller left CM in comparison to control group participants (t(63) = -2.62, p = 0.028). The same pattern was observed in left Pf nucleus (t(62) = -2.58, p = 0.031). More pronounced statistical differences were found upon comparison of GMV in right CM and Pf nuclei in patients compared to controls (F(2,83) = 5.2, p = 0.007, 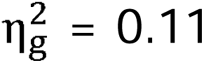 and F(2,82) = 4.25, p = 0.018, 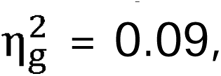 respectively). Group with left-sided facial pain showed no GMV reduction in CM and Pf (p > 0.05). Patients and healthy participants also differed in sizes of paratenial nuclei (F(2,83) = 5.32, p = 0.007, 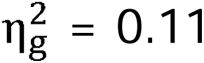). Specifically, grey matter volume of left paratenial nucleus was lower in patients with right-sided pain than in healthy control group (t(63) = -3.47, p = 0.002). Group of patients with left-sided pain did not show any statistically significant difference neither with right-sided pain group nor with healthy controls (p > 0.05). The size of right Pt was different across groups (F(2,83) = 4.4, p = 0.015, 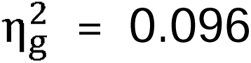). In contrast to the left Pt, GMV of right paratenial nucleus differed significantly not only between group with right-sided pain and controls, but also between controls and left-sided pain group (t(63) = -2.48, p = 0.04 and t(49) = -2.46, p = 0.042, respectively).

LGN and MGN nuclei concerned with visual and auditory signal processing, respectively, underwent the most pronounced GMV reductions in the PTN patients. Individuals with either side of pain demonstrated substantial alterations in both right and left lateral geniculate bodies (F(2,82) = 8.58, p < 0.001, 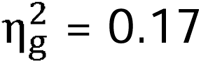 and F(2,82) = 10.9, p < 0.001, 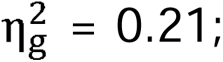 Fig.3 A and B). Patients with right-sided pain had lower values of GMV in LGN bilaterally (t(63) = -3.94, p < 0.001 for right LGN and t(63) = -4.63, p < 0.001 for left LGN). The same trend was evident in patients suffering left-sided pain (t(49) = -3.06, p = 0.008 for right LGN and t(49) = -2.7, p = 0.023 for left LGN). The volume of the left medial geniculate body varied between groups as well (F(2,83) = 8.45, p < 0.001, 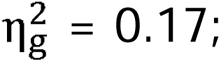 Fig. 3 C). Both patients with left- and right-sided pain had significant differences from control group in measured GMV (t(50) = -3.24, p = 0.005 and t(63) = -3.74, p < 0.001, respectively). However, we did not find any differences in the volume of right MGN between groups (p > 0.05). Among the ventral group of nuclei processing somatosensory signals, we were able to discover only between-group differences in GMV of left ventral lateral posterior (VLp) and ventral posterolateral (VPL) nuclei (F(2,81) = 3.6, p = 0.032, 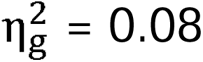 and F(2,82) = 3.98, p = 0.022, 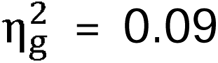). These nuclei showed a pronounced decrease in volume in patients having right-sided pain (VLp: t(61) = -3.1, p = 0.007; VPL: t(63) = -2.77, p = 0.019).

**Figure 3.**
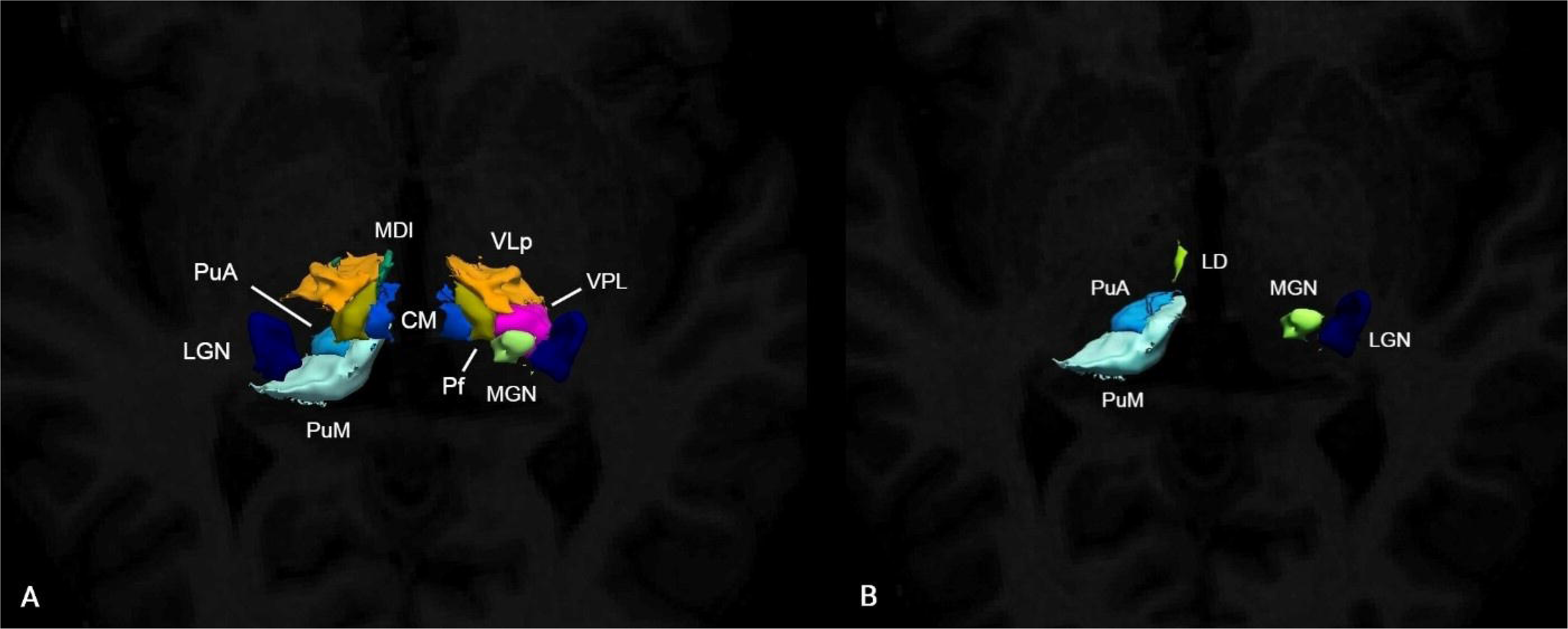
Panel A. Reconstruction of thalamic nuclei showing significant differences between patients with right-sided pain and healthy controls. Panel B. Visualization of nuclei whose volumes differed significantly between patients with left-sided pain and control group’s individuals. In both cases the size of depicted nuclei was larger in healthy participants. Paratenial nuclei are not shown due to their small size. CM - centromedian nucleus, LGN - lateral geniculate nucleus, MDl - mediodorsal lateral nucleus, MGN - medial geniculate nucleus, Pf - parafascicular nucleus, PuA - anterior pulvinar nucleus, PuM - medial pulvinar nucleus, VLp - ventral lateral posterior nucleus, VPL - ventral posterolateral nucleus.

**Figure 4.**
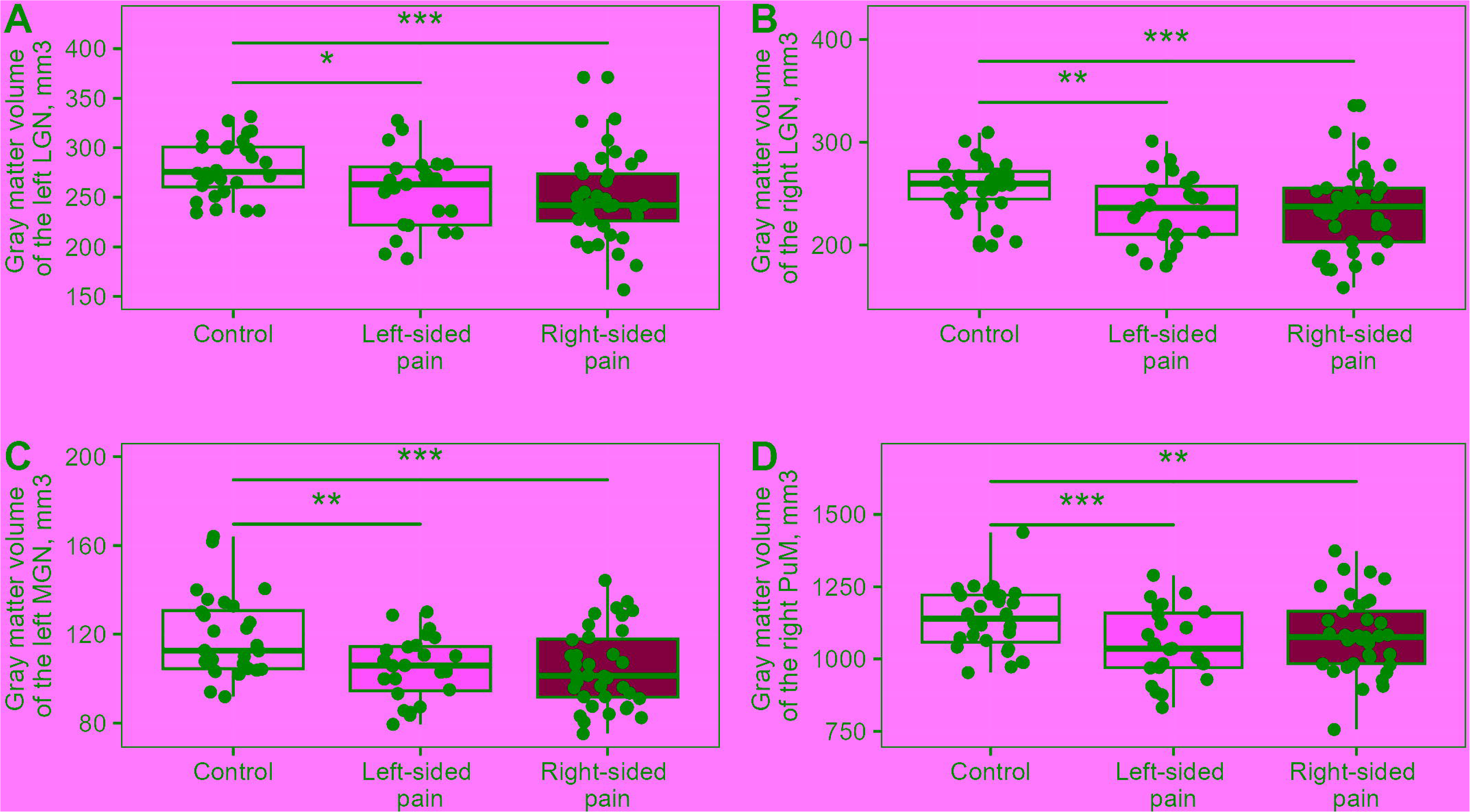
The figure depicts box plots showing the distribution of data in each of the experimental groups for thalamic nuclei with the largest effect size. Groups demonstrating statistically significant differences in mean gray matter volumes are indicated by lines with asterisks, reflecting the level of statistical significance.

In addition, right, but not left anterior (PuA) and medial pulvinar (PuM) regions were also characterized by GMV reduction in PTN patients (F(2,82) = 4.08, p = 0.02, 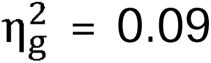 and F(2,82) = 8.43, p < 0.001, 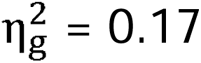). Patients of both groups had smaller volumes of right PuA compared to healthy subjects (t(49) = -2.46, p = 0.042 for patients with left-sided pain and t(63) = -2.48, p = 0.04 for patients with right-sided pain). Similarly, right medial pulvinar underwent profound reduction of gray matter in patients with either side of pain (left: t(49) = -3.9, p < 0.001; right: t(63) = -3.04, p = 0.009; Fig 3 D). Left pulvinar regions in patients did not show any differences from the control group’s ones (p > 0.05).

Volume reduction was also evident in associative right LD and MDl nuclei (F(2,83) = 3.69, p = 0.029, 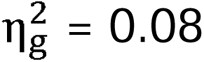 and F(2,83) = 3.89, p = 0.024, 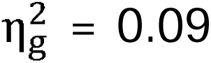). Mean values of LD volume were distinct in patients with left-sided pain and controls (t(50) = -2.6, p = 0.029), whereas MDl sizes were shown to be different between controls and patients with right-sided pain (t(63) = -2.79, p = 0.018). Differences in GMV of individual thalamic nuclei between patients with right- and left-sided pain were not statistically significant for all above-mentioned comparisons (p > 0.05). The visualization of thalamic nuclei, which exhibited statistically significant differences compared to the control group, for patients with right-sided and left-sided pain is presented in Figure 2. The results of the comparison between a single group of patients with trigeminal neuralgia, without stratification based on the presence of right- or left-sided pain, and healthy controls are provided in the Supplementary file.

### Follow-up MRI study

MRI data of 34 patients collected 6 months after surgery was used for the follow-up morphometric analysis of thalamic nuclei. To ensure reliability of results, we first checked whether all differences detected between groups at the preceding stage were retained on a subsample of smaller size (n = 34). Volumes of left Pf (W = 605, p = 0.11) and right LD (W = 616, p = 0.13), MDl (W = 636, p = 0.15), PuA (W = 620, p = 0.07), VLp (W = 624, p = 0.13) were found to be statistically indistinguishable from those of controls. Among the nuclei that did differ between groups, none displayed an increase in GMV 6 months after surgical intervention (p > 0.05). We conducted additional analysis by excluding 5 patients who did not demonstrate 100% pain reduction after the surgery and thus were classified as non-responders according to our preset criteria. No changes were observed compared to the results of the previous analysis. Detailed information on all statistical comparisons is available in the Supplementary file.

### Gray matter volume assessment in patient groups undergoing different surgical procedures

Fifty-six patients from our sample underwent surgical intervention. Specifically, forty patients were subjected to microvascular decompression surgery (MVD; 64%), 10 patients underwent radiofrequency rhizotomy (RF; 16%) and balloon compression was performed on 6 patients (BC; 10%). Of these patients, 45 individuals (80%) were considered responders meaning that they were fully pain-free for at least six months post-surgery. 11 patients (20%) either did not respond to surgical treatment or surgery did not lead to full pain relief and demonstrated only partial effects. The gray matter volume of thalamic nuclei did not differ among patients in groups with favorable and unfavorable outcomes of surgical interventions (p > 0.05). For the purpose of statistical analysis, patients undergoing radiofrequency rhizotomy and balloon compression were combined into one group (percutaneous interventions, PI). The other two groups comprised healthy subjects and patients who underwent microvascular decompression.

In most cases, statistically significant differences in gray matter volumes of thalamic nuclei between groups arose from distinctions between healthy subjects and patients undergoing microvascular decompression (see Table 4 for specific values). The most pronounced differences between these groups were shown in the left LGN (p < 0.001), left LP nucleus (p = 0.001), right CM (p = 0.001) and right LGN (p < 0.001).

**Table 4.**
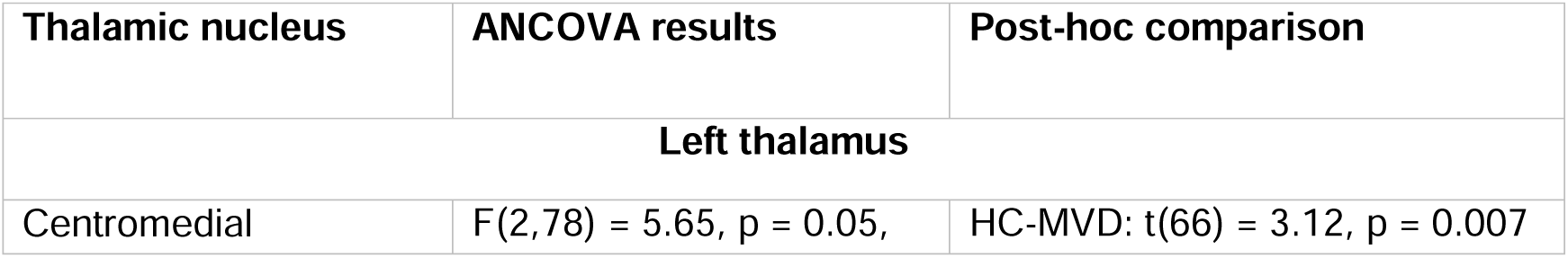
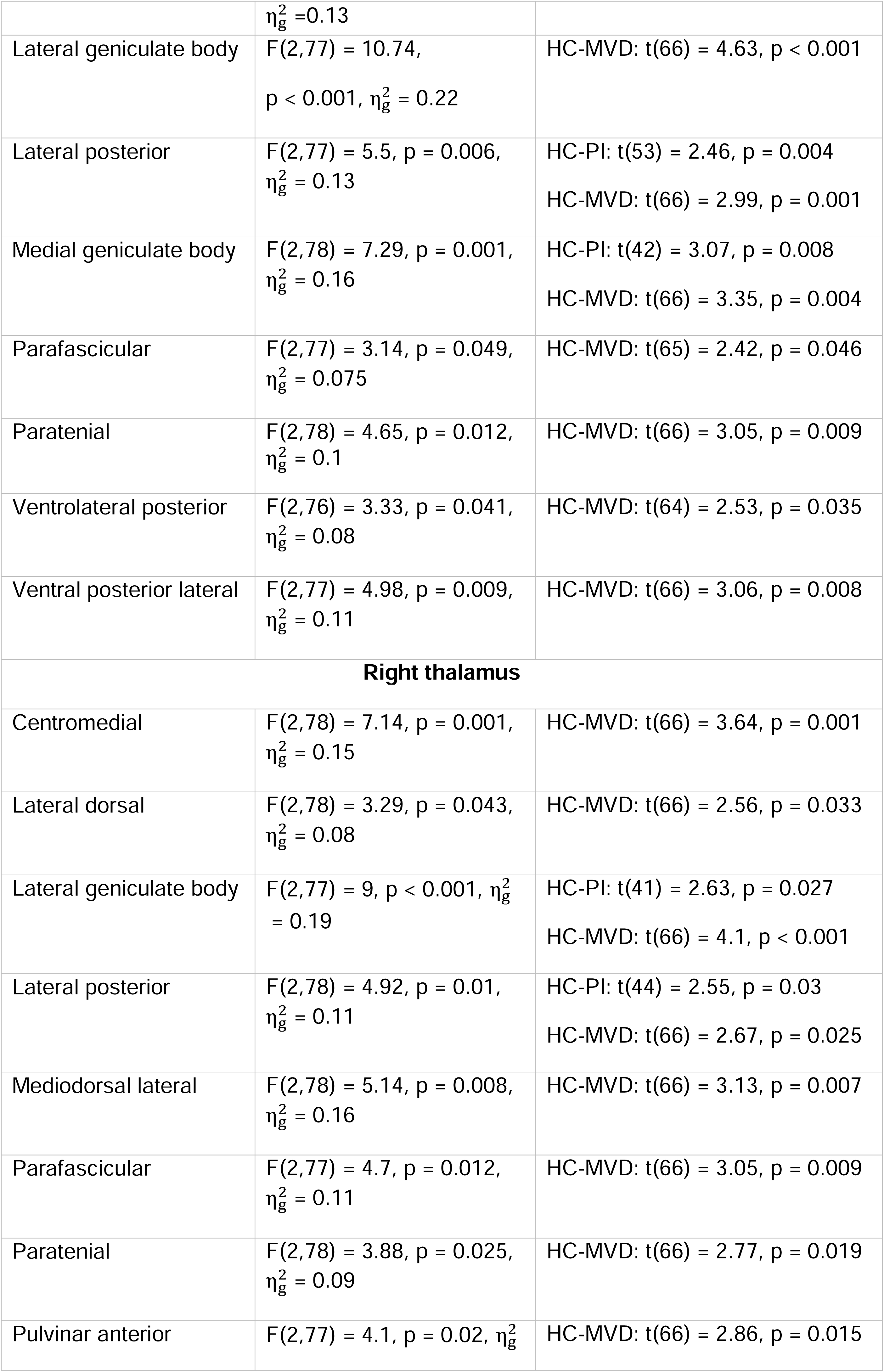
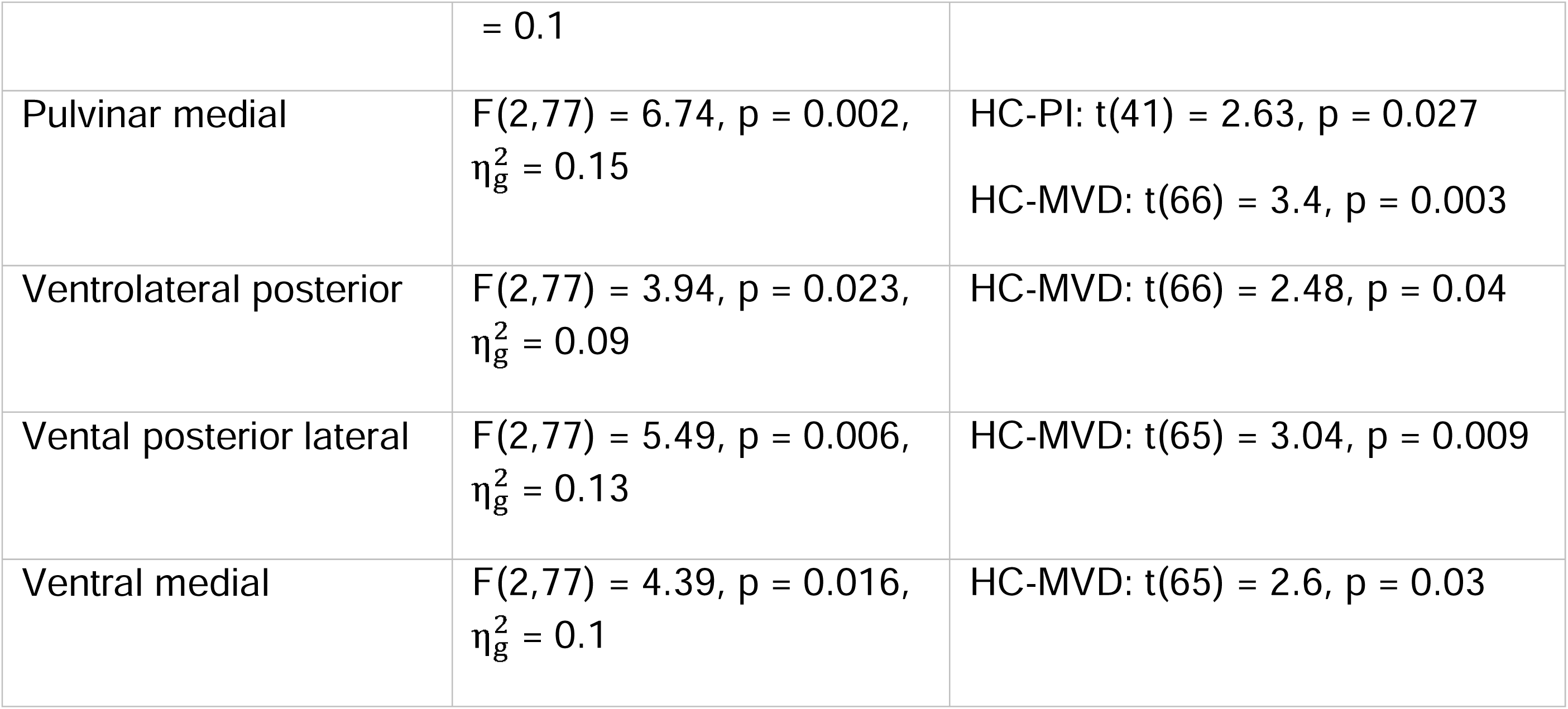
The gray matter volume of thalamic nuclei, demonstrating statistically significant differences between groups of healthy participants and patients undergoing microvascular decompression or percutaneous interventions.

Patients undergoing percutaneous interventions had lower values of GMV compared to HC group in left lateral posterior nucleus (p = 0.004) and medial geniculate body (p = 0.008), as well as in right lateral posterior nucleus (p = 0.03), medial pulvinar (p = 0.027) and lateral geniculate body (p = 0.027), which were also accompanied by distinctions between HC and MVD groups.

### Associations between morphometric and clinical variables

We ran a correlational analysis to uncover relations between the extent of structural alteration of individual thalamic nuclei and a number of clinical variables like disease duration and pain intensity (taken as mean BPI score). It is noteworthy that all significant correlations that survived FDR correction were found only for patients with right-sided pain, and subsequently, only these correlations are presented in this section. Disease duration did not show any significant correlations with thalamic GMV (all p > 0.1). In contrast, pain intensity was negatively correlated with GMV in the right/left LGN (r = - 0.58, p = 0.009, CI95 = [-0.78, -0.28]; r = -0.56, p = 0.01, CI95 = [-0.76, -0.25], respectively), right/left Pt (r = -0.51, p = 0.019, CI95 = [-0.74, -0.19]; r = -0.46, p = 0.03, CI95 = [-0.71, -0.13]), bilateral VM (r = -0.58, p = 0.009, CI95 = [-0.78, -0.28] for right VM and r = -0.48, p =0.025, CI95 = [-0.71, -0.14] for left VM), bilateral VPL (r = -0.55, p = 0.013, CI = [-0.76, -0.23] for right VPL and r = - 0.5, p = 0.019, CI95 = [-0.73, -0.18] for left VPL). Additionally, we found strong negative correlations between right anterior, inferior and medial pulvinar size and pain intensity (r = - 0.48, p = 0.025, CI95 = [-0.71, -0.15]; r = -0.67, p = 0.001, CI95 = [-0.83, -0.42]; r = -0.56, p = 0.01, CI95 = [-0.76, -0.25], respectively. The only left-sided pulvinar nucleus that was correlated with the pain intensity was the medial one (r = -0.49, p = 0.025, CI95 = [-0.72, - 0.16]).

Additionally, we sought to characterize the influence of drug intake on measured variables. Specifically, for this analysis, we selected only those patients who were taking carbamazepine as a sole medicine. We found that the dose of carbamazepine did not correlate significantly with GMV in thalamic nuclei and patient’s main clinical variables (p > 0.05).

**Figure 5.**
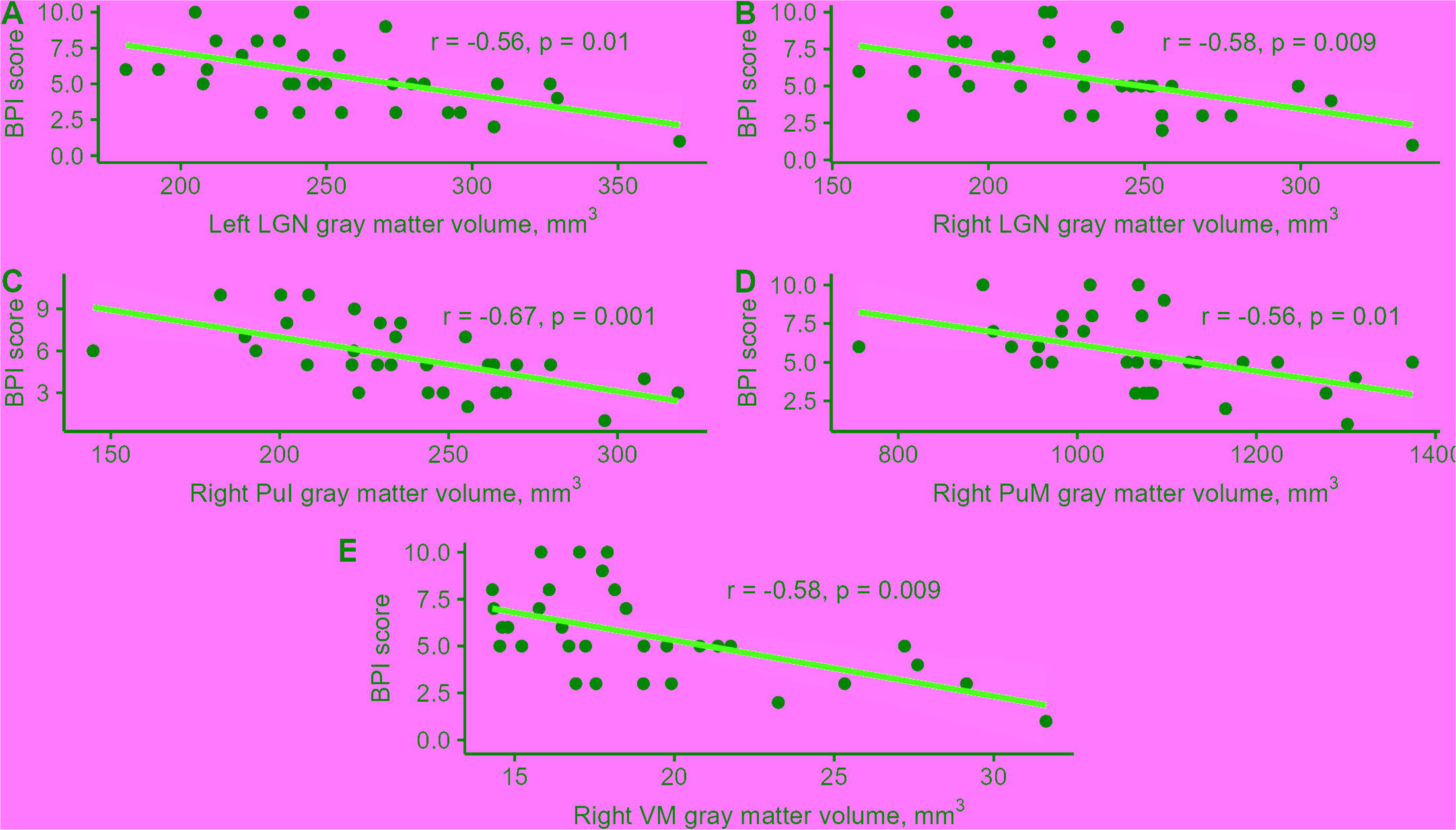
The correlation plots displaying relationship between pain intensity and gray matter volumes in the thalamic nuclei in patients with right-sided facial pain. A. Left LGN. B. Right LGN. C. Right PuI. D. Right PuM. E. Right VM. The red lines represent a linear fit to the data points depicted in black. The shaded areas around these lines correspond to 95% confidence intervals.

## Discussion

Structural remodeling of thalamic nuclei has been shown to take place in a wide range of brain disorders [14]. Results of our previous study showed fractional anisotropy decrease in thalami of patients with primary TN [15]. In this study, we for the first time investigated thalamic nuclei in patients with chronic TN and demonstrated that they underwent significant structural alterations over the course of disease. Besides, we were able to estimate the relationship between volumes of individual thalamic nuclei and a number of clinical and neuropsychological variables. In doing so, we found that CM and Pf nuclei, being an integral part of the intralaminar group, LGN, MGN, VPL, which belong to the sensory cluster, VLp involved in the motor processing and associative PuA and PuM demonstrated the most noticeable alterations in their gray matter volumes.

CM-Pf complex, along with some nuclei of ventral group, including VLp, represents one of the major thalamic recipients of abundant afferents originating in basal ganglia and has long been known to be an effective target of deep brain stimulation in patients with intractable movement disorders and disorders of consciousness [26,44]. It has also previously been shown in a number of early and more recent papers that CM-Pf stimulation can help to successively mitigate intense chronic pain in some patients [1]. Alternatively, a stereotactic radiosurgery technique, medial thalamotomy, targeting CM-Pf also holds promise due to its pain-alleviating effects [16]. Importantly, the CM-Pf nuclei extensively innervate primary and secondary cortical motor areas. Transcranial stimulation of the motor cortex has emerged as another safe and effective option for managing otherwise untreatable pain, as supported by a series of clinical trials conducted in the past decade [34]. Anterior pulvinar, which projects widely to sensorimotor cortical areas and medial frontal gyrus was also shown to be structurally altered in our patients and may be involved not only in the action planning and executing domains, but also in higher-order cognitive functions [4,19]. Furthermore, it is now recognized that post-stroke lesions of anterior pulvinar frequently manifest themselves in the form of intense pain [40]. The involvement of brain motor areas observed in patients suffering chronic pain may stem from the fact that they are often forced to restrict their movements and learn to act differently in order not to provoke or exacerbate pain. Consistent with this trend, studies focusing on patients with trigeminal neuralgia have also reported noticeable structural changes occurring in motor-related areas of these individuals [12, 36].

Unlike the currently emerging interest in studying the motor side of pain experience, the sensory component of pain has always been a main focus of investigation in individuals with chronic pain. Earlier studies have consistently demonstrated the GMV changes in brain regions involved in nociceptive processing, including thalamic nuclei [42]. In line with these findings, our own results point to the reduction of gray matter volume in ventral posterolateral nucleus (VPL), which is believed to be engaged in processing of incoming nociceptive afferents and relaying them to upstream somatosensory cortical regions. This finding can be interpreted as a compensatory strategy employed to cope with a significantly augmented influx of pain signals [33]. In addition, we were able to detect even more pronounced differences in volumes of lateral and medial geniculate bodies between groups. To the best of our knowledge, this is the first direct evidence for deep neural reorganization that takes place at the level of sensory thalamus in patients with PTN. The shift in this subtle balance between visual, auditory and nociceptive processing in case of chronic pain might be driven by redistribution of priority given to each of the sensory modalities due to the heightened salience of pain signals. The indirect corroboration of this hypothesis comes from studies demonstrating that CM-Pf, which exhibited shrinkage in our patients, might also be implicated in attention switching and reorienting toward salient objects or scenes [6,32]. However, in our study all thalamic nuclei that differed between PTN and HC groups were reduced in volume in a sample of patients and we did not find any conclusive evidence for the enlargement of nuclei involved in nociceptive processing.

Since pain is a complex biological phenomenon commonly defined as having tightly interwoven sensory, affective and cognitive components, another viable alternative may be that this reduction of LGN and MGN is secondary to alterations of brain areas concerned with affective processing. For instance, prior studies by Apkarian and colleagues convincingly demonstrated preferential recruitment of limbic regions as opposed to somatosensory areas during fMRI sessions in patients with chronic pain [3]. Moreover, these regions were also subject to considerable gray matter volume loss [21, 41]. There is also substantial evidence indicating a high prevalence of anxiety and depression among patients suffering from chronic trigeminal neuralgia [9]. Recent discoveries have highlighted the significant structural and functional changes that can occur in visual and auditory brain areas, which are directly connected to the LGN and MGN, as a result of comorbid conditions [10, 31]. Notably, the medial pulvinar, which exhibits gray matter volume reduction in patients with trigeminal neuralgia, is particularly implicated in various functional domains associated with emotional processing. It is known to receive dense afferents from the amygdala, hippocampus, posterior insula, anterior and posterior cingulate cortex, as well as from extensive regions of the frontal and parietal lobes [35]. Lesions involving the medial pulvinar commonly manifest as selective or more generalized impairments in emotional recognition and expression [7, 43]. Recent studies have further demonstrated the involvement of the medial pulvinar in regulating arousal, which varied in response to nociceptive stimulation [5]. Interestingly, surgical intervention known as pulvinotomy has been found to result in a noticeable reduction in pain intensity and improved quality of life [45].

In this study, we were also able to find that patients with different pain sides diverged in terms of the number of thalamic nuclei showing marked differences from controls. Volume reduction was restricted to paratenial, laterodorsal, anterior and medial pulvinar, as well as lateral and medial geniculate nuclei in individuals with left-sided pain. In contrast, patients with right-sided pain were characterized by an additional drop of GMV in CM-Pf, MDl, VPL. LD size was not significantly different from that of healthy participants. Henssen and colleagues previously hypothesized an existence of an ipsilateral pathway for transmission of nociceptive information, alongside the well-known contralateral one [20]. Overall, a flux of nociceptive afferentation may not be symmetric for right and left sides of the body and strongly influenced by top-down regulation systems, including descending pain inhibitory system [39]. Overexcitation induced by an intensified stream of nociceptive signals might be a driving factor for gray matter loss in a range of thalamic nuclei. In addition, future studies are needed to ensure that these results are not just a consequence of the different number of patients in each of the groups.

In our sample, pain intensity and disease duration displayed negative correlations with volumes of nuclei. It is particularly noteworthy that the decrease of GMV in all thalamic nuclei was not reversed 6 months after surgical intervention implying that prolonged experience of pain may lead to a drastic and long-standing remodeling of established patterns of neural interactions. These findings also prompt the consideration of distinguishing between PTN-related structural changes and preexisting reorganized neural circuitries that may serve as risk factors for the transition from acute to chronic pain in patients with predispositions.

Patients with chronic pain commonly showcase neuropsychological deficits across a number of domains [2,8]. We investigated the extent to which these domains were affected in patients with trigeminal neuralgia and found that only verbal fluency scores differed between experimental groups. Moreover, they were positively correlated with gray matter volumes in some affected thalamic nuclei, although the strength of these correlations was generally moderate at best. Verbal fluency performance heavily relies on executive functions, which are frequently impaired in chronic pain and can contribute to less favorable treatment outcomes [13,22]. Further investigations are necessary to comprehensively explore the complex bidirectional relationship between pain and cognition.

Taken together, these findings have promising clinical implications for uncovering previously unknown details regarding chronic pain development and guiding clinical decision-making. However, it is worthwhile to mention several limitations of this study. First, we were unable to detect a statistically significant impact of carbamazepine intake on thalamic nuclei gray matter volume and cognitive functions in patients, despite subjective reports of memory decline and difficulties in attention focusing by the patients. This issue should be further studied systematically and in greater depth in future research. Second, current study design does not allow us to rule out the possibility that observed structural alterations in thalamic nuclei were secondary to that of in the basal ganglia or cortex. Future prospective longitudinal studies are required to address this issue. Finally, several investigated nuclei that showed statistically significant differences between groups during pre-surgical assessment, did not considerably differ at the 6 months follow-up on a smaller sample, which implies that caution should be taken when using these results to inform clinical decisions.

## Authorship contribution statement

AP: conceptualization, methodology, data collection, formal analysis, writing - original draft, reviewing & editing. EF: conceptualization, methodology, data collection, formal analysis, reviewing & editing. BZ: data collection and formal analysis. AM: data collection. GM: conceptualization, methodology, reviewing and editing, supervision, project administration. JR: conceptualization, methodology, reviewing and editing, supervision, project administration. All authors had editorial input to the manuscript.

## Data statement

all raw data is available from the corresponding author upon reasonable request.

## Data availability statement

Data available on request from the authors

