## Supplementary files for "Thalamic nuclei in patients with chronic facial pain: gray matter volume patterns before and after surgery"

Neurocognitive assessment

MMSE scale is intended to serve as a brief and easily administered tool for cognitive screening, enabling prompt detection of severe cognitive impairments. In the current study, patients who scored less than 24 (out of 30) points on this test were excluded from the further analysis.

Rey–Osterrieth complex figure task was carried out in three parts. In the first part participants are asked to copy the given complex figure as precisely as possible. Second part is an immediate recall phase, wherein the patient’s aim is to draw the earlier shown figure from memory. The same task given to patients twenty minutes later constituted a delayed recall phase. The final scoring was conducted according to the scheme introduced by Petilli and colleagues and involved dividing the complex figure into 48 basic elements, awarding 1 point for each correctly reproduced element ([Petilli et al., 2021](https://www.nature.com/articles/s41598-021-94247-9)).

Trail making test consisting of two main parts is one of the most widely used and reliable tools for evaluating visual attention and behavioral flexibility in clinical practice. In the part A, the subject is given instruction to connect a series of 25 dots in an ascending numerical order with the goal of completing the task as rapidly as possible while ensuring accuracy is maintained. In its turn, part B places an extra emphasis on attention flexibility, complicating the task with an addition of the requirement of switching between digits and letters in the following way: 1-A, 2-B, 3-C e t.c. Since direct TMT scores, that is, parts A and B completion times, are strongly affected by age, education and other sociocultural factors, in this study, ratio score (B/A) was used as an attention performance index instead ([Christidi et al., 2015](https://pubmed.ncbi.nlm.nih.gov/25798536/)).

Digit Span is a cognitive assessment tool used to measure verbal short-term and working memory. It consists of two formats: forward digit span (DS-F) and backward digit span (DS-B). In this verbal task, auditory stimuli are presented to the participants, who are then required to verbally repeat the series of digits either in the same order as presented (forward span) or in reverse order (backward span). The FDS assesses verbal working memory and attention, while the BDS additionally evaluates cognitive control and executive functions.

Verbal fluency tests are another commonly utilized measure to assess executive dysfunction and typically involve generating words within a specific timeframe. Two main types of these tasks were employed in this study, namely, phonological verbal fluency task, where words starting with a given letter are produced, and categorical verbal fluency task, where words within a specific semantic category are generated. The final score for each task was a number of words participants could generate in 60 seconds. These tasks measure the functioning of the semantic system, which is responsible for storing words and their associations. Additionally, they evaluate the efficacy of retrieval strategies and the individual's capacity for self-monitoring and inhibiting inappropriate responses.

|  | **Responders** | **Non-responders** |
| --- | --- | --- |
| MVD | 33 | 7 |
| RF | 7 | 3 |
| BC | 5 | 1 |

**Supplementary Table 1.** Frequencies of responders and non-responders across surgery types

| **Branches affected** |  |
| --- | --- |
| V1 | 13 (21%) |
| V2 | 54 (87%) |
| V3 | 39 (63%) |

| **Surgery type** |  |
| --- | --- |
| MVD | 40 (64.5%) |
| Radiofrequency rhizotomy | 10 (16.1%) |
| Balloon compression | 6 (9.7%) |
| No surgery | 6 (9.7%) |
| **Surgery outcome** |  |
| Responders | 45 (80.4%) |
| Non-responders | 11 (19.6%) |
| **Complications** |  |
| No complications | 38 (67.9%) |
| Minor | 11 (19.6%) |
| Major | 7 (12.5%) |
| **Previous surgeries** | 14 (23%) |

**Supplementary Table 2.** Clinical data of patients.

| **Thalamic nucleus** | **Wilcoxon test statistics** |
| --- | --- |
| Left CM | W = 366, p = 0.25 |
| Left LGN | W = 242, p = 0.5 |
| Left MGN | W = 340, p = 0.48 |
| Left Pf * | W = 414, p = 0.05 |
| Left Pt | W = 385, p = 0.14 |
| Left VLp | W = 382, p = 0.1 |
| Left VPL | W = 248, p = 0.57 |
| Right CM | W = 270, p = 0.65 |
| Right LD * | W = 433, p = 0.058 |
| Right LGN | W = 280, p = 1 |
| Right MDl | W = 375, p = 0.19 |
| Right MGN | W = 321, p = 0.7 |
| Right Pf | W = 397, p = 0.06 |
| Right Pt | W = 398, p = 0.09 |
| Right PuA | W = 223, p = 0.31 |
| Right PuM | W = 257, p = 0.68 |
| Right VLp | W = 399, p = 0.13 |

**Supplementary Table 3.** Results of MRI follow-up study (before surgery and 6 months after).

| Thalamic nucleus (HC-PTN comparison) | p-value (FDR-corrected) |
| --- | --- |
| Left AV | 0.47 |
| Left CeM | 0.79 |
| Left CL | 0.46 |
| Left CM | 0.1 |
| Left LD | 0.1 |
| Left LGN | 0.00075 |
| Left LP | 0.1 |
| Left L-Sg | 0.15 |
| Left MDl | 0.27 |
| Left MDm | 0.79 |
| Left MGN | 0.0012 |
| Left MV(Re) | 0.27 |
| Left Pc | 0.27 |
| Left Pf | 0.14 |
| Left Pt | 0.057 |
| Left PuA | 0.43 |
| Left PuI | 0.45 |
| Left PuL | 0.93 |
| Left PuM | 0.22 |
| Left VA | 0.4 |
| Left VAmc | 0.45 |
| Left VLa | 0.15 |
| Left VLp | 0.1 |
| Left VM | 0.31 |
| Left VPL | 0.085 |
| Right AV | 0.38 |
| Right CeM | 0.38 |
| Right CL | 0.16 |
| Right CM | 0.024 |
| Right LD | 0.049 |
| Right LGN | 0.002 |
| Right LP | 0.14 |
| Right L-Sg | 0.1 |
| Right MDl | 0.03 |
| Right MDm | 0.14 |
| Right MGN | 0.049 |
| Right MV(Re) | 0.26 |
| Right Pc | 0.049 |
| Right Pf | 0.029 |
| Right Pt | 0.029 |
| Right PuA | 0.029 |
| Right PuI | 0.085 |
| Right PuL | 0.8 |
| Right PuM | 0.0048 |
| Right VA | 0.1 |
| Right VAmc | 0.09 |
| Right VLa | 0.1 |
| Right VLp | 0.09 |
| Right VM | 0.14 |
| Right VPL | 0.06 |

**Supplementary Table 4.** Results of comparison of thalamic nuclei GMV between groups
